# Meta-aligner: Long-read alignment based on genome statistics

**DOI:** 10.1101/060129

**Authors:** Damoun Nashta-ali, Ali Aliyari, Mohammad Amin Edrisi, Ahmad Ahmadian Moghadam, Seyed Abolfazl Motahari, Babak Hossein Khalaj

## Abstract

Fast and accurate alignment of long-reads plays an important role in reducing the overall cost of long-read sequencing. In this paper, we propose Meta-aligner, an efficient and accurate long-read aligner that exploits the statistics of reference genome to improve performance in terms of reducing time complexity and achieving significantly higher recall for very noisy and long reads. The first step of algorithm adopts well-known short-read aligners in order to rapidly align a large fraction of reads through a progressive process of aligning read fragments to the reference genome. In the second phase, the remaining reads are handled by simultaneous alignment of all read fragments and a decision making process which exploits the overall information provided by the corresponding mapped fragments. By using this procedure, significant performance improvement is attained in comparison with traditional schemes in the case of PacBio long-reads.

---

The number of short and long reads produced by Next Generation Sequencing (NGS) technologies is growing very rapidly. Evidently, efficient and accurate mapping of these reads to the reference genome plays an important role in reducing the overall NGS cost as well as improving downstream analysis in applications such as re-sequencing, RNA-Seq, and ChIP-Seq. Currently, NGS technologies can be divided into two categories based on the overall quality and length of the reads. Sequencers such as Illumina-HIseq and Ion Torrent-Proton with short and almost clean reads fall in the first category, while PacBio-RS II and Nanopore-MinION are typical examples of sequencers that provide long but noisier reads.

A number of algorithms and softwares, such as Bowtie [1], mrsFAST [2], SOAP2 [3] are typically used for alignment of short reads, while Bowtie2 [4], BWA [5], and Seqalto [6] are among methods used for handling long reads. Comparative analysis of these aligners is not trivial as quality can be measured in terms of a number of metrics such as handling reads within the repeat regions, handling very noisy reads, uniqueness and accuracy in the report, speed, and finally, the effect of results on a given downstream analysis [7].

In this paper, we are concerned with alignment of long and very long reads (longer than 300 base pairs). One popular design technique in this regime is to extract *seeds* (small fragments) from the reads and find exact or very similar matches for these seeds within the reference genome. After anchoring extracted seeds to several locations, local alignment algorithms are used to determine the best match(es) for the given read. In general, one common theme among the available long-read aligners is that they treat all reads equally. However, as shown in this paper, differentiating between reads that are coming from repeat and non-repeat regions leads to significant improvement in performance of the alignment scheme. In fact, the key contrast between our approach and traditional aligners is that we focus on exploiting the inherent genome structure and the underlying statistics of the reads from the outset.

From a high level perspective, Meta-aligner consists of two different stages, namely alignment and assignment. The first stage is designed to rapidly and accurately align easily mappable reads to the reference genome using traditional short-read aligners for mapping small fragments of reads uniquely. As our results clearly demonstrate, due to statistical properties of real genomes, a large fraction of reads will be handled at this stage. The remaining reads are relatively harder to align and, therefore, additional processing at the assignment stage should be devoted to properly align them. These reads are handled by aligning all remaining small fragments of them. However, as the number of reads processed at the second stage is relatively small, the overall time complexity of the second stage is less than the first stage.

## RESULTS

### Genome’s Statistics: random intervals

As an example, consider the case of resequencing problem where reads are mapped to the reference genome and variants are called from mapped reads. In the ideal case, the goal is to map all reads uniquely to their original positions in the reference genome. However, the repeat structure of genomes makes it difficult to map all the reads correctly and uniquely to the genome.

Our goal is to show that by taking into account the genome structure, a large number of reads can be aligned very rapidly and accurately. To this end, we first consider a model in which the reference genome resembles a conceptual mosaic structure alternating between two types of intervals, namely repeat and random intervals (as shown in Supplementary 1 Fig. 5). The type of interval can be defined based on two parameters *ℓ* and *d*. A repeat (respectively random) interval is defined as consecutive bases where any substring of length *ℓ* starting from a position within the interval can be matched to some other (respectively no other) location(s) of the genome with maximum Hamming distance *d*. Thus, any substring of length *ℓ* is classified as either a repeat or a random interval. Supplementary 1 Figure 2 shows how the fraction of random intervals changes as *ℓ* increases from 20 to 200 for *d* = 0 in chr19 of hg19. This figure shows that about 98 percent are categorized as non-repeat regions for the choice of *ℓ* = 80 with *d* = 0.

In the next step, we extend the mosaic model of the reference genome to the reads themselves, as every read also consists of mosaic random and repeat intervals. In fact, the mosaic structure of the read is exactly copied from its originating position on the reference genome. In this way, the *ℓ* spectrum of the read can be partitioned into random and repeat *ℓ*-mers where a random/repeat *ℓ*-mer is defined as an *ℓ*-mer located within a random/repeat interval of the original genome. It should be highlighted that, in fact, the *ℓ*-mers are the ones that play the key role in the read mapping process as they can be mapped uniquely to the genome within a maximum Hamming distance 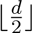.

Our proposed algorithm rests on the aforementioned model of the genome and exploits it to map reads from a target genome correctly and uniquely to the reference genome. Let us consider an algorithm that attempts to uniquely map all the *ℓ*-mers (the *ℓ*-spectrum) of a given read with maximum Hamming distance 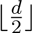 to the reference genome. We also assume that such algorithm anchors the read to the locations where at least one *ℓ*-mer is mapped uniquely. In this way, the algorithm outputs a list of possible locations for each read. We are interested in the case where only one consistent interval exists in the genome where *all* the uniquely mapped *ℓ*-mers of a given read belong to; this location will then be identified as the true position of that read. However, two types of reads do not map uniquely to their true positions: (a) reads that none of their *ℓ*-mers can be mapped uniquely to the reference genome, i.e. the whole read resides inside the repeat interval or (b) reads which at least one of their *ℓ*-mers is uniquely mapped to an incorrect location. In the following, we will address handling of the first and, then, the second of aforementioned read types through an additional decision making procedure.

It is important to note that the fraction of repeat intervals as defined in our model, in addition to the number of anchored reads are both functions of parameters such as *ℓ*, *d*, and read length *L*. In order to investigate the effect of these parameters on the number of reads of the first type, let us consider the statistics of the number of reads with uniquely mappable *ℓ*-mers in chr19 of hg19. It is expected that an increase in *ℓ* results in an increase in the number of random *ℓ*-mers within a read as shown in Figure 1. In this simulation, we scan the whole chr19 and generate all reads of length *L* that start at every base. However, the number of reads without any random *ℓ*-mer surprisingly remains almost constant for different values of *ℓ* and *d* in a typical range considered in our simulations.

As shown in Figure 1(a) and (c), only ≈ 0.49% and ≈ 1.18% of reads of length *L* = 1,000 bps have no random *ℓ*-mer (first type of reads) of length *ℓ* = 40 for *d* = 0 and *d* = 3, respectively. The same pattern is also observed for the choice of smaller read length of *L* = 400 bps. It should also be noted that although aligning all the *ℓ*-mers to the reference genome increases the sensitivity of the algorithm, it also incurs additional computational complexity that should be taken into account. For instance, such full *ℓ*-mer alignment requires mapping 960 40-mers to anchor a read of length 1,000 with *ℓ* = 40. One way to significantly decrease the order of complexity is to consider only non-overlapping *ℓ*-mers. As such sampling, in general, leads to some level of performance degradation, we have reported the results in the case of non-overlapping *ℓ*-mers in Table (a). The results show negligible change in the number of reads of the first type when such non-overlapping *ℓ*-mers are used. In fact, only ≈ 0.54% and ≈ 2.26% of reads of length *L* = 1, 000 bps have no non-overlapping random *ℓ*-mer of length *ℓ* = 40 for *d* = 0 and *d* = 3, respectively. However, in the case of shorter reads, the number of non-overlapping *ℓ*-mers does not provide enough diversity for a given read and the chance of having at least one random *ℓ*-mer reduces. For example, for *L =*400 bps, ≈ 0.7% and ≈ 6.3% of reads have no non-overlapping random *ℓ*-mer of length *ℓ* = *40* for *d* = 0 and *d* = 3, respectively. In such cases, it is suggested to use a number of overlapping *ℓ*-mers (using a sliding window of length Si) to increase diversity.

**Table I.** Analysis of random and low-repeat *ℓ*-mers. (a) Percentage of disjoint random *ℓ*-mers within reads of lengths *L* = 400 and *L* = 1000 of ch19 of hg19. (b) and (c) Fraction of the remaining reads after the first step and their number of low-repeat *ℓ*-mers with different list sizes *ℒ*_*s*,1_ = {5,10, 20,40} for *ℓ* = 40. In (b) and (c), we assume that, all t-mers and only non-overlapping *ℓ*-mers, are respectively used at the first step.

| | Number of disjoint random $\ell$-mers within a read<br>for $L = 400$ | | | | Number of disjoint random $\ell$-mers within a read<br>for $L = 1000$ | | | |
| --- | --- | --- | --- | --- | --- | --- | --- | --- |
| | 0 | 1 | between 1 and $\approx L/\ell$ | $\approx L/\ell$ | 0 | 1 | between 1 and $\approx L/\ell$ | $\approx L/\ell$ |
| $(d, \ell) = (0, 20)$ | **0.9%** | 0.7% | 69.1% | 29% | **0.6%** | 0.19% | 85.6% | 13.5% |
| $(d, \ell) = (0, 40)$ | **0.7%** | 0.4% | 39.4% | 59.1% | **0.54%** | 0.17% | 62.6% | 36.68% |
| $(d, \ell) = (\mathbf{3}, \mathbf{40})$ | **6.3%** | 8% | 84.8% | 0.8% | **2.26%** | 1.49% | 96.2% | 0.04% |
| $(d, \ell) = (0, 80)$ | **0.7%** | 0.3% | 4.2% | 94.8% | **0.49%** | 0.17% | 8.2% | 91.1% |

(a)
| $\mathcal{L}_{s,1}$ | Number of low-repeat $\ell$-mer within a read<br>for $d=0$ | | | Number of low-repeat $\ell$-mer within a read<br>for $d=3$ | | |
| --- | --- | --- | --- | --- | --- | --- |
| | 0 | between 1 and 80 | between 81 and $\approx L/\ell$ | 0 | between 1 and 80 | between 81 and $\approx L/\ell$ |
| 5 | 56.73% | 11.94% | 30.98% | 32.01% | 30.97% | 36.88% |
| 10 | 54.46% | 4.9% | 4.9% | 26.21% | 19.81% | 53.83% |
| 20 | 52.75% | 0.08% | 46.81% | 21.90% | 4.06% | 73.89% |
| 40 | 52.75% | 0.08% | 46.81% | 21.69% | 0.07% | 78.09% |

(b)
| $\mathcal{L}_{s,1}$ | Number of low-repeat $\ell$-mer within a read<br>for $d=0$ | | | Number of low-repeat $\ell$-mer within a read<br>for $d=3$ | | |
| --- | --- | --- | --- | --- | --- | --- |
| | 0 | between 1 and 80 | between 81 and $\approx L/\ell$ | 0 | between 1 and 80 | between 81 and $\approx L/\ell$ |
| 5 | 52.31% | 12.28% | 35.08% | 17.18% | 20.15% | 62.59% |
| 10 | 50.22% | 4.91% | 44.54% | 14.07% | 11.67% | 74.18% |
| 20 | 48.64% | 0.1% | 50.93% | 11.75% | 2.24% | 85.93% |
| 40 | 48.64% | 0.1% | 50.93% | 11.64% | 0.04% | 88.23% |

In the next step, we address the case of second type of reads with at least one of their *ℓ*-mers uniquely mapped to an incorrect location. In practice, we are interested in handling very noisy reads where substitutions and indels are inevitable. In such scenarios, some *ℓ*-mers are still uniquely, but now incorrectly, mapped to the reference genome. In order to address this issue, two different approaches can be adopted: (a) decision making based on more than one uniquely mapped *ℓ*-mers that correspond to the same location or (b) using longer *ℓ*-mers to alleviate the effect of noisy reads. For example, in the case of 40-mers, one may consider mapping either two different 40-mers or single 80-mers. The first approach is more favorable, since anchoring longer *ℓ*-mers is time consuming especially in case of high error rates. Additionally, addressing indels and structural variations in case of longer *ℓ*-mers leads to a more challenging gap-based alignment step. In fact, shorter *ℓ*-mers can be aligned with or without small gaps and it may still be possible to find enough *ℓ*-mers that are not significantly exposed to structural variations and/or indels. Finally, repeats can be bridged by two or more *ℓ*-mers similar to mate-pairs. In genomes with high fraction of repetitive areas, such as human genome, the second approach anchors the reads efficiently and correctly to the repeat intervals if the flanking parts of the read still falls in a random interval.

In order to maximize the throughput of the algorithm, it is important to properly choose the values of *ℓ* and *d* for a given reference genome, sequencing error rate, and variation rate. In Supplementary 1 Section 1, assuming an
i.i.d. reference and error model, we analytically determine the appropriate values of *ℓ* and *d* for different sequencing error and variation rates. We show that *ℓ* = 40 is the proper choice of *ℓ* for such model, when sequencing error rate is lower than 10%. Subsequently, the value of *d* is obtained by computing the average number of errors within the substring of length *ℓ* = 40. As we will show in the sequel, our method can additionally exploit the real genome structure, by adopting an estimation mechanism (Parameter Estimation algorithm) that can find the proper values of parameters *ℓ* and *d* from the given reads and corresponding reference genome. The details of such estimation mechanism and its results are provided in Supplementary 1 Section 3 and Supplementary 2.

### Genome’s Statistics: repeats with low copy numbers

Using uniquely mapped *ℓ*-mers, most of the reads are uniquely and correctly mapped to the reference genome, with a remarkably low computational complexity. The remaining reads which are not anchored are mostly reads that come from pseudogenes and gene families that are classified as low copy number repeats. However, anchoring these reads is still highly valuable for downstream genome analysis. In order to handle this class of reads, the proposed algorithm maps all *ℓ*-mers with maximum Hamming distance 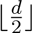 to the reference genome and reports a finite list of size *ℒ*_*s*,1_ of possible positions for each *ℓ*-mer. All *ℓ*-mers whose number of possible positions exceeds the list size *ℒ*_*s*,1_ are discarded, and the *ℓ*-mer to which a list of size *ℒ*_*s*,1_ greater than one is assigned is labeled as a *low-repeat ℓ*-mer. The reads that we are considering at this stage fall into two categories: (a) for the first type of reads, none of their *ℓ*-mers are low-repeat *ℓ*-mer, in which case that read will be discarded, or (b) for the second type of reads, the list assigned to at least one of their low-repeat *ℓ*-mers does not include its correct location. In the next step, we will first discuss what percentage of reads are of the first type. Subsequently, we will present a method for identifying correct possible positions for the second type of reads.

Let us assume that in the previous part, all *ℓ*-mers of each read are used for anchoring that read and after this part a number of reads are still remaining to be further processed. Table (b) shows the percentage of remaining reads of length *L* = 1,000 bps as a function of number of low-repeat 40-mers. In the case of *ℒ*_*s*,1_ = 10 and *d* = 0, only ≈ 54.46% of the remaining reads have no low-repeat 40-mers (≈ 0.27% of total reads) and changing *d* to *d* = 3 has negligible effect on those percentages. Thus, the first type of reads constitute only a small fraction of reads that can then be aligned using larger *ℓ*-mers and list sizes.

Table (c) shows the results of same process when only non-overlapping *ℓ*-mers of reads were used through the same process. In this case, with the same *ℒ*_*s*,1_ = 10 and *d* = 0, ≈ 50.22% of the remaining reads have no low-repeat *ℓ*-mers (≈ 0.27% of total reads). These results show that in the case of long enough reads, using only the non-overlapping *ℓ*-mers has almost the same performance as using all *ℓ*-spectra.

In order to handle the second type of reads for which the list corresponding to some low-repeat *ℓ*-mers does not contain their correct positions, we look for maximum consensus positions among the lists. Details are presented in the Online Methods.

### Real data and synthetic simulation results

In the current implementation of Meta-aligner, the following short-read aligners can be used in the alignment stage: Bowtie [1], SOAP2 [3], and mrsFast-Ultra [2], where Bowtie is set as the default aligner. In Supplementary 1 Section 4, we simulated the alignment stage of Meta-aligner for these three short-read aligners and show that Bowtie is the best aligner due to its performance. In the assignment stage, Bowtie and Bowtie2 are used to create a list of candidate positions for small and long fragments, respectively. We use sub-fragment lengths *ℓ*_1_ at the first and second steps and *ℓ*_2_ at the third step of Meta-aligner, respectively. The overall structure of Meta-aligner is shown in Supplementary 1 Figure 7.

All simulations are executed on 24 threads of a 24-core cluster with 32 GB RAM. The human genome hg19 including sex chromosomes is used as the reference genome. We have simulated Meta-aligner as well as several long-read aligners (Seqalto, Bowtie2 and BWA) to align a real dataset. The real dataset is from Human 54x PacBio reads with accession number SRX533609 published in NCBI GenBank ”http://www.ncbi.nlm.nih.gov/sra?term=SRX533609”, where the first two files, SRR1304331 and SRR1304332 were used. We selected 174,537 reads of this dataset with minimum and average lengths of 500 and 6890, respectively, without any constraint on their qualities (see Supplementary 1 Section 8). Applying PE algorithm to this dataset results in two sets of parameters in terms of trade-off between run time and mapping rate: 1) when run time has a higher priority: (*ℓ*_1_,*d*) = (25,1) and no sliding, 2) when favoring mapping rate: (*ℓ*_1_,*d*) = (25,2) with *𝒮*_1_ = 5. The PE algorithm also estimates the mismatch and indel error rates of the PacBio read set as, 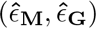 = (**2.8**%, **13.8**%).

Seqalto and Bowtie2 failed to align this read set due to high error rates and RAM constraints (even at 192 GB RAM), respectively. Comparison of Meta-aligner and BWA-SW results is presented in Table II. BWA-SW with default parameters (*z* = 2) has smaller run time, but results in lower number of aligned reads compared to Meta-aligner. The mapping rate of BWA-SW can be improved by setting *z* = 10 at the expense of higher time complexity. The important fact is that since PacBio reads are noisy and dominated by indels, BWA-SW shows inferior performance in the case of unique alignment as it reports many candidate locations for each read and the reported locations are usually clumped close to each other, leading to confusion for down-stream analysis. In overall, BWA-SW uniquely aligns 21% and 19.61% of reads with *z* = 2 and *z* = 10, respectively. However, Meta-aligner aligns 87.04% of reads uniquely at the first stage and 93.37% of reads at the end of the second step. For more details, see Supplementary 1 Section 7.

**Table II.** PacBio read set simulation results.

| Aligner | Commands or Parameters | Mapping (%) | Time(sec) | Peak Memory (GB) |
| --- | --- | --- | --- | --- |
| Meta-aligner | $(\ell_1, d, S_1) = (25, 1, -)$ | 88.99 | 24,160 | 27 |
| Meta-aligner | $(\ell_1, d, S_1) = (25, 2, 5)$ | 97.97 | 39,041 | 27 |
| BWA-SW | $z = 2$ | 90.46 | 1,573 | 22.6 |
| BWA-SW | $z = 10$ | 95.55 | 207,838 | 28.5 |

In order to investigate robustness of Meta-aligner to variations in read lengths and quality scores, the number of aligned reads of Meta-aligner in three different scenarios are shown in Figure. In this figure, the number of input reads are shown in red as a reference.

Figure (a) shows that almost all reads longer than 4 Kbps are uniquely aligned after the alignment stage of Meta-aligner. Subsequently, the assignment stage significantly increases the number of aligned reads for reads shorter than 4 Kbps. Figure (b) shows that all reads with quality scores above 0.84 are aligned at the end of both stages of Meta-aligner. For lower quality reads, only a small fraction of reads are remained unaligned. For instance, 84.4% of the ≈ 3,200 reads with quality scores lower than 0.7 are handled by Meta-aligner after the second step.

In order to highlight the strength of Meta-aligner in the case of both noisy and long reads, we generated *N* synthetic reads using an i.i.d. error model with *ϵ* = {10; 15; 20}%, and read lengths *L* = {1,000; 4,000; 8,000} bps, from hg19 where *N* ⨯ *L* = 10^9^. Table III shows performance of Meta-aligner for this scenario. In fact, at high indel rates, Metal-aligner shows better performance as its design inherently takes such issues into account.

In order to evaluate the performance of Meta-aligner in the case of shorter reads, we generated *N* = 1, 000, 000 synthetic reads using an i.i.d. error model with *ϵ* = {2; 5; 10} % and read lengths *L* = {300; 500; 1,000} bps, from the hg19. Our comparative results, as shown in Figure demonstrate that at the given error rates and read lengths, Meta-aligner’s performance in terms of recall rate and precision remained at very high quality levels. As expected, Meta-Aligner’s performance improves as read length is increased. Although BWA-SW takes less run time, its recall and precision are worse than Meta-aligner. In the case of Seqalto, it failed to align any reads with *L* = 1,000 bps and *ϵ* = 10%. In addition, its run time for *L* = 1,000 and e = 5% is over 100,000 seconds, which is out of range of Figure. Fraction of unique reports for these simulations is also provided in Supplementary 1 Figures 10, 11. These results show that large fraction of reads are uniquely reported for Meta-aligner (in comparison with other aligners). More results are presented in Supplementary 1 Section 6.

## DISCUSSION

We propose Meta-aligner, a new method for alignment of long-reads based on the genome structure. We show that genome structure has implicit information which should be taken into account for designing a more efficient alignment algorithm. Meta-aligner handles reads classified based on the genome structure within three classes. First, reads with many short sub-fragments originating from unique regions of the genome are aligned uniquely, fast and almost accurately by only considering two sub-fragments of such read. The remaining reads are categorized into two classes based on the number of copies of their short sub-fragments as either low or high copy number reads. Reads for which a major number of sub-fragments come from low copy number regions of the genome are handled by assigning a small list to them. Subsequently, the same procedure is executed for high copy number reads with a larger list. Results show the accuracy and high mapping percentage of Meta-aligner for long PacBio reads. Another key point to note is that Meta-aligner has a mechanism to estimate its key parameters from the input read set and reference genome to further optimize its performance for real genomes by adopting to their statistics.

**Table III.**
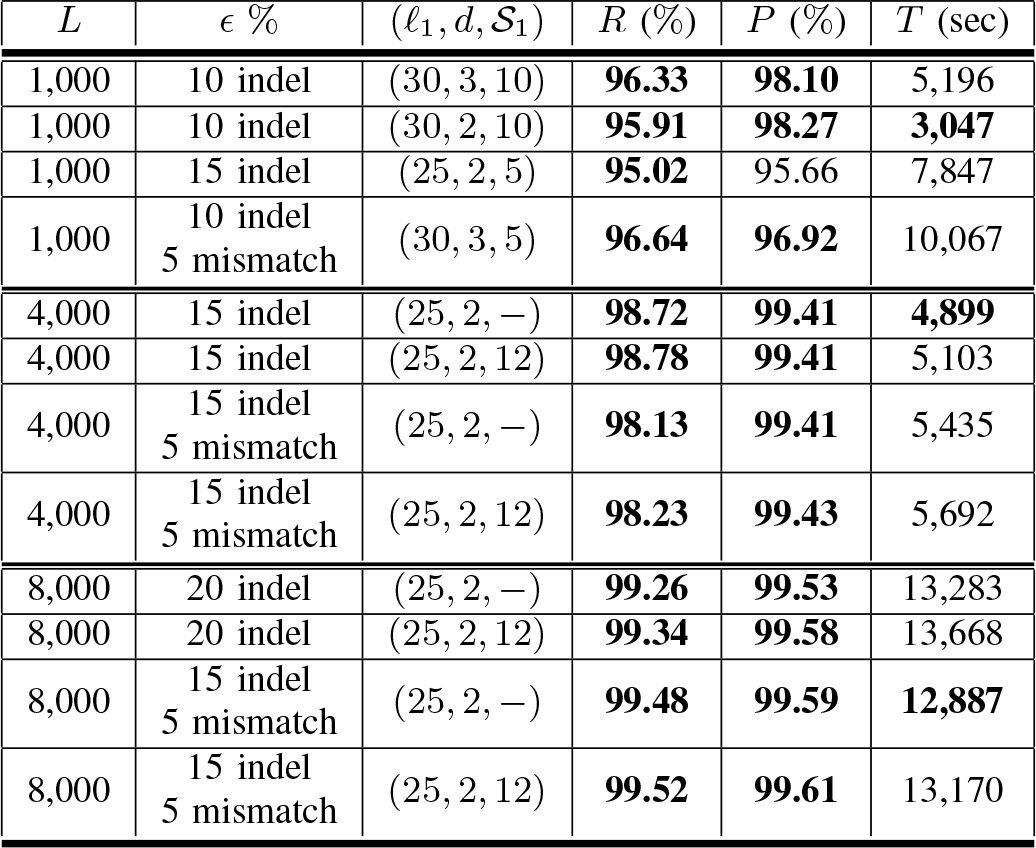
The overall Meta-aligner results for hg19 in the case of very high error rates (*ϵ* = {10; 15; 20} %) and read lengths of *L* = {1, 000; 4, 000; 8, 000} bps). Number of reads of length *L* is *N* = 10^9^/*L*.

## METHODS

Methods and any associated references are available in the online version of the paper at online method.

**Figure 1.**
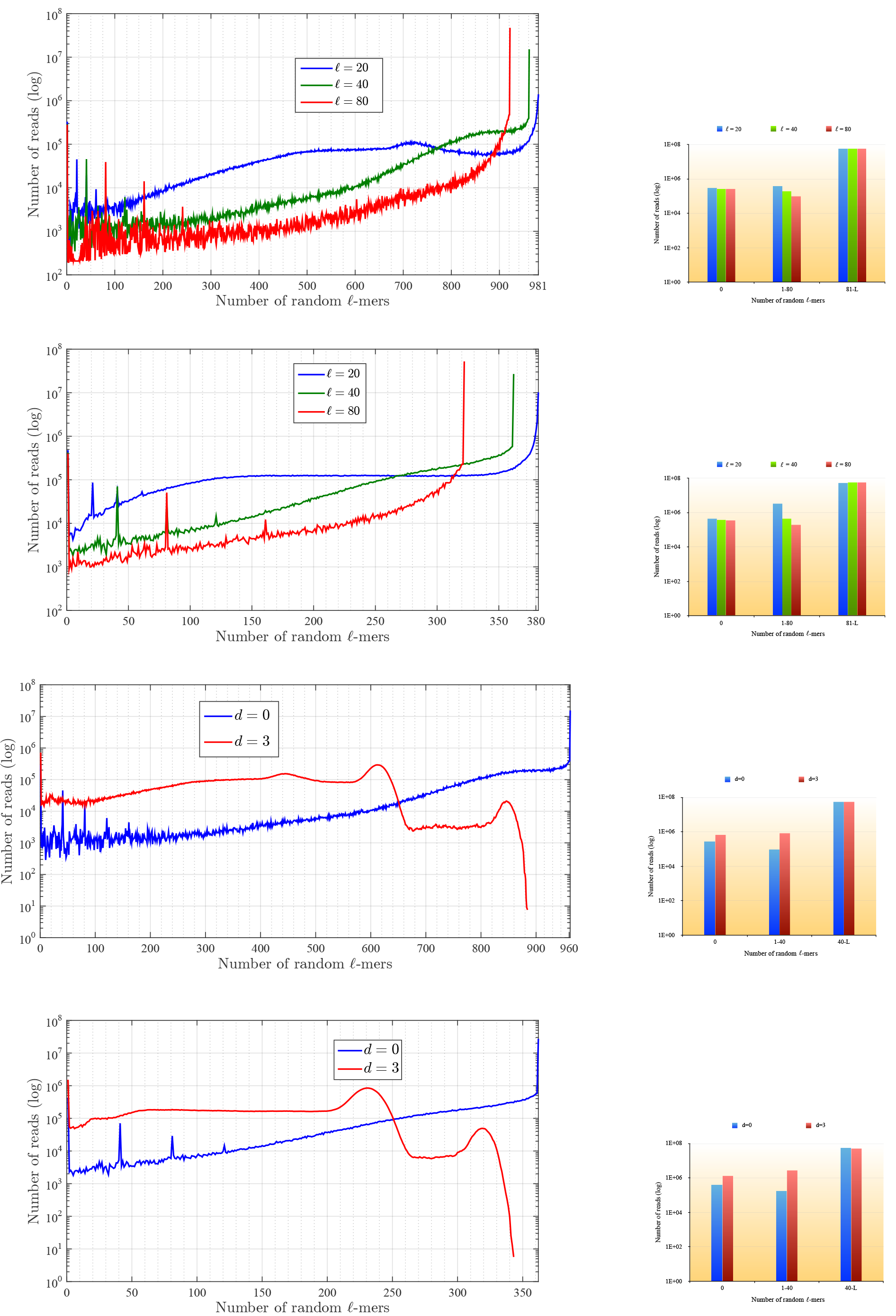
Distribution of random t-mers within reads in chr19 of hg19. (a) Number of reads of length 1,000 with random *ℓ*-mers for different values of *ℓ* and *d* = 0. The right figure shows the histogram of reads with 0, [1-80] and [81-(*L* - *ℓ*)] random *ℓ*-mers. (b) Number of reads of length 400 with random *ℓ*-mers for different values of *ℓ* and *d* = 0. The right figure shows the histogram of reads with 0, [1-80] and [81-(*L* — *ℓ*)] random *ℓ*-mers. (c) Number of reads of length 1,000 with random t-mers for different

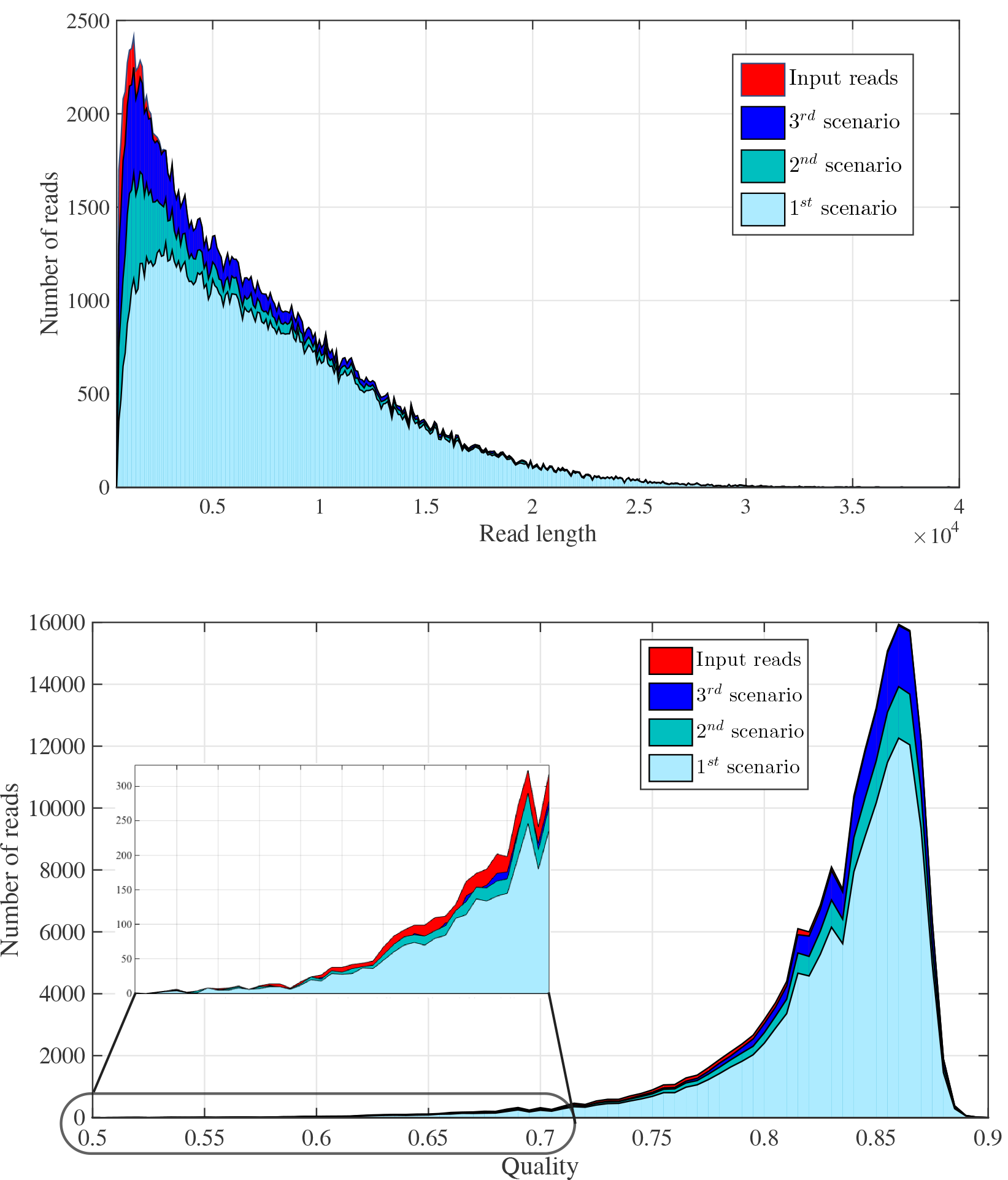

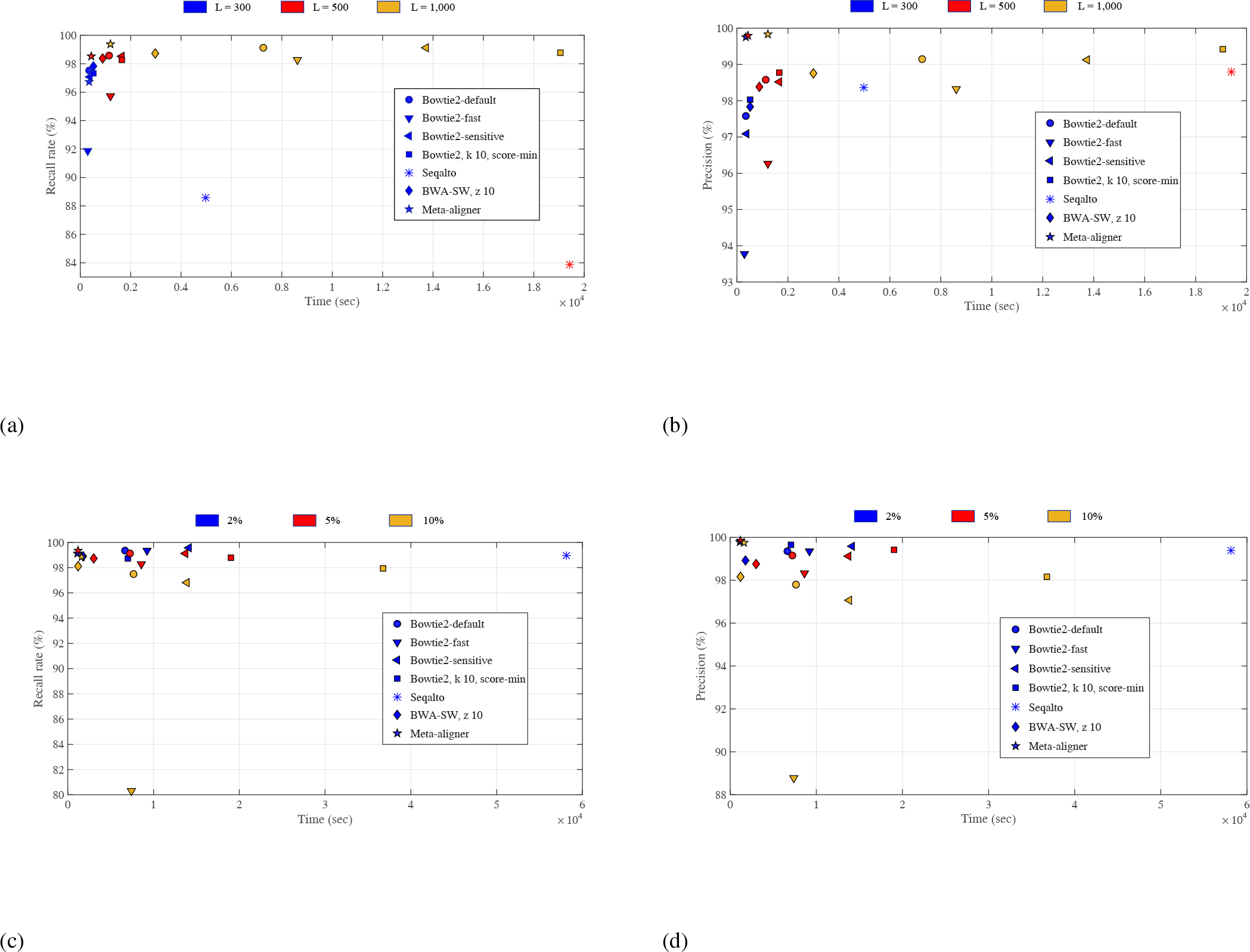

## ONLINE METHOD

**Parameters definition and synthetic read set construction**: The variations between target and reference genomes is statistically modelled by randomly dispersed substitutions and indels. It is also assumed that the variation rate is *α* The reads are also assumed to be noisy with an error rate ϵ which is a mixture of substitutions and indels. Let *ℛ* denote the set of sequenced reads of length *L*. Meta-aligner method consists of a PE algorithm for parameters estimation and two stages, namely *alignment* and *assignment.* The proposed PE algorithm, alignment and assignment procedures are illustrated in Supplementary 1 Figures 7-9 and are detailed in Supplementary 1 Algorithms 1-4.

**PE algorithm**: In the PE algorithm, Meta-aligner estimates the values *ℓ*, *d*, mismatch and indel error rates (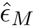 and 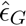), and the normalized cutting distance for local alignment. These parameters are estimated by running alignment stage for the first *N_t_* input reads. Meta-aligner first aligns these reads with *ℓ* = 25 and *d* = 2. Then, local alignment of the anchored reads provides an estimation of mismatch and indel error rates. Subsequently, Meta-aligner estimates the best pair (*ℓ*, *d* = 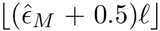) for values of *ℓ ϵ* {15,20, 25,30,35, 40,45, 50} to achieve the lowest anchoring time. Subsequently, its recall rate is close to the maximum recall rate of this set. The normalized cutting distance for local alignment is set to 5 ⨯ 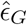.

**Anchoring step of the alignment stage**: In the alignment stage, all reads are first divided into non-overlapping consecutive smaller fragments of length *ℓ*_1_ forming an array of *K_i_* = ⌊*L*_*i*_/*ℓ*_1_⌋ fragments for the *i*^th^ read of length *L_i_*. The 1^st^ and 2^nd^ fragments of all reads are aligned to the reference genome using any aligner capable of mapping reads with a Hamming distance of *d*_1_. If any of these fragments is uniquely aligned, a tag is assigned to the fragment, representing its corresponding position and flag (forward or reverse directions) on the reference genome. In the case that two fragments are uniquely aligned and they *confirm* each other, that read is anchored to the confirmed position. The first two fragments *confirm* each other if the difference of their aligned positions is in the range

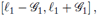

and have the same flags. Parameter 𝒢_1_ is set to accommodate for indels within two fragments of length *ℓ*_1_. Reads with confirmed fragments are removed from the set of reads *ℛ*. Subsequently, the third fragment of the remaining reads are aligned to the reference genome, and the reads are anchored if two fragments out of three confirm each other. The anchored reads are then removed from the set. As explained in Supplementary 1 Algorithm1, the fragments are progressively added and anchored reads with exactly two confirmed fragments are removed from the set. In general, the *i*^th^ and *j*^th^ uniquely aligned fragments of each read *confirm* each other if the difference of the mapped positions satisfies

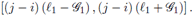

The procedure stops whenever *ℛ* = *Ø* or all fragments are processed. Up to this step, we have considered nonoverlapping fragments. However, the aforementioned algorithm can be applied to overlapping fragments as well. For this case, anchoring is repeated ⌈*ℓ*_1_/*𝒮*_1_⌉ times, such that at the *i*^th^ step, we start from the (*i* - 1) ⨯ 𝒮_1_-th base of all reads, for all *i* ∈ {1, …, ⌈ℓ_1_/𝒮_1_⌉}. We recommend that overlapped fragments are used for very noisy read and reads of length less than 300.

**Local alignment step of the alignment stage**: The anchored reads can be aligned by any local alignment algorithm. We use Smith-Waterman, which provides the highest sensitivity. Such high sensitivity is necessary for long-reads with high error rate. We consider the normalized cutting distance which shows the number of columns and rows that is used in the local alignment table relative to each read length, in the local alignment process. This parameter is between 0 and 2 and its default value is 0.05.

**Constructing tables of the assignment stage**: At this stage, all reads remaining in *ℛ* are divided into overlapping fragments of length *ℓ*_1_. A default value for the overlapping fragments’ length is initially set to *ℓ*_1_/2. All fragments are then aligned by considering a maximum list of size *ℒ*_*s*,1_ consisting of possible positions with a distance smaller than *d*_1_ from the reference genome. Subsequently, fragments with more than ℒ_*s*_,_*i*_ positions are discarded. The list of positions of the *i*^th^ fragment of a given read *r* is denoted by *T*_*i*_(*r*). In order to emphasise the number of reported positions for each fragment, we assign a score to each reported position as a function of the reported aligner score and its number of reported positions. We propose an exponential weight to incorporate the effect of number of reported positions, as follows,

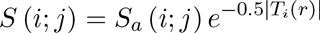

where *S_a_* (*i*; *j*) and |*T*_*i*_(*r*)| denote the reported score of the aligner for the *j*^th^ element of *T_*i*_* and the size of *T_i_* (*r*), respectively.

**Constructing paths of the assignment stage**: Consider a position *p ∈ T*_1_ (*r*). If *p* confirms another position *q ∈ Tj* (*r*), ∀*j* > 1, we construct a *path* consisting of *p* and *q*, subsequently removing them from the tables. We continue this procedure by searching for all possible paths consisting of the remaining positions within tables. The score of each path is set to the sum of scores of all its positions. Subsequently, paths are sorted in a decreasing order. In most cases, a gap separates the paths with relatively higher scores from those with relatively smaller scores. The paths with relatively higher scores are then selected according to the parameter (𝒮_th_) which determines the threshold of relatively higher scores.

**Selecting paths of the assignment stage**: After local alignment of each path of *r* to the corresponding location of the genome, paths with low alignment scores are filtered out and a list of possible positions is reported for *r*. Reads with any reported list are removed from *ℛ*.

**Repeating the assignment stage**: After this step, the same procedure can be applied to all reads in *ℛ* with larger fragments of length *ℓ*_2_ > *ℓ*_1_ and larger maximum list size of ℒ_2_.

## AUTHOR’S CONTRIBUTIONS

DN, SAM performed the analysis and discussed the results of Meta-aligner. AA, AAM, and MAE were mainly responsible for implementing Meta-aligner with help from DN. DN, SAM and BHK wrote the manuscript.

## ACKNOWLEDGEMENTS

We would like to thank Mostafa Tavassolipour for his valuable helps.

## ADDITIONAL FILES

*Additional file 1- Supplementary 1 file*

*Additional file 1- Supplementary 2 file*

